# The di-symbiotic systems in the aphids *Sipha maydis* and *Peryphillus lyropictus* provide a contrasting picture of recent co-obligate nutritional endosymbiosis in aphids

**DOI:** 10.1101/2022.06.12.495806

**Authors:** François Renoz, Jérôme Ambroise, Bertrand Bearzatto, Samir Fakhour, Nicolas Parisot, Mélanie Ribeiro Lopes, Jean-Luc Gala, Federica Calevro, Thierry Hance

**Author notes:** Address correspondence to François Renoz.

## Abstract

Dependence on multiple nutritional bacterial symbionts forming a metabolic unit has repeatedly evolved in many insect species that feed on nutritionally unbalanced diets such as plant sap. This is the case for aphids of the subfamilies Lachninae and Chaitophorinae, which have evolved di-symbiotic systems in which the ancient obligate nutritional symbiont *Buchnera aphidicola* is metabolically complemented by an additional nutritional symbiont acquired more recently. Deciphering how different symbionts integrate both metabolically and anatomically in such systems is crucial to understanding how complex nutritional symbiotic systems function and evolve. In this study, we sequenced and analyzed the genomes of the symbionts *B. aphidicola* and *Serratia symbiotica* associated with the Chaitophorinae aphids *Sipha maydis* and *Periphyllus lyropictus*. Our results show that, in these two species, *B. aphidicola* and *S. symbiotica* complement each other metabolically (and their hosts) for the biosynthesis of essential amino acids and vitamins but with distinct metabolic reactions supported by each symbiont depending on the host species. Furthermore, the *S. symbiotica* symbiont associated with *S. maydis* appears to be strictly compartmentalized into the specialized host cells housing symbionts in aphids, the bacteriocytes, whereas the *S. symbiotica* symbiont associated with *P. lyropictus* exhibits a highly invasive phenotype, presumably because it is capable of expressing a larger set of virulence factors, including a complete flagellum for bacterial motility. Such contrasting levels of metabolic and anatomical integration for two *S. symbiotica* symbionts that were recently acquired as nutritional co-obligate partners reflect distinct coevolutionary processes specific to each association.

## Introduction

Many insect species depend on inherited endosymbiotic bacteria to access certain nutrients and have evolved specialized cells - the bacteriocytes - to host these nutritional partners [1]. Bacteriocytes are the interface for metabolic exchanges between the host and the symbionts. They allow the host to control symbiont populations according to its nutritional needs and ensure stable vertical transmission of symbionts from mother to offspring [2,3]. This compartmentalization into bacteriocytes is considered as the outcome of a long coevolutionary history between the insect hosts and their obligate bacterial partners [4]. However, a consequence of this intracellular lifestyle is the isolation of symbionts from any environmental source of prokaryotic DNA, leading to Muller’s ratchet, i.e. the accumulation of irreversible deleterious mutations in the symbiont genome [5]. Combined with severe population bottlenecks during vertical transmission and the relaxation of purifying selection on genes that have become unnecessary in a context of interdependent association, this leads to the evolutionary degeneration of the symbiont genome [6,7]. Thus, extreme genomic reduction is a typical feature of obligate insect nutritional symbionts, often acquired tens of millions years ago [8,9]. This degeneration, once far advanced, can jeopardize the symbiotic functions of the nutritional symbiont and ultimately represents an evolutionary dead end for host lineages [10–12].

An evolutionary solution to this impasse is the acquisition by the host species of additional symbionts that evolve as co-obligate nutritional partners by forming a metabolic unit with the ancestral obligate symbiont that has become unable to perform its nutritional function alone [13]. The existence of such multi-symbiotic nutritional systems has been reported in various sap-feeding insect species [13–22] with evidence showing that the acquisition of additional obligate symbionts is a very dynamic process involving recurrent recruitments of new bacterial symbionts and repeated replacements of pre-existing intracellular ones [17,19,23,24]. While the nutritional basis of co-obligate symbioses has been examined in different sap-feeding insects through genomic analyses [13], many aspects of the biology of these associations remain unexplored, including the infection mechanisms that the co-obligate symbionts may use to colonize host tissues and the localization of these bacteria within the host. Examining these aspects in light of the diversity of co-obligate nutritional endosymbiosis is critical for understanding the evolution, development, and functioning of multi-symbiotic nutritional systems in insects.

Aphids (Hemiptera: Aphididae) of the subfamilies Lachninae and Chaitophorinae are valuable models for addressing the biology of co-obligate symbiosis in insects. In these aphids, the eroded metabolic abilities of the ancient obligate symbiont *Buchnera aphidicola* are complemented by those of a more recently acquired co-obligate symbiont, often belonging to the *Serratia symbiotica* species, one of the most common secondary symbionts found in aphids. Specifically, *B. aphidicola* and *S. symbiotica* metabolically complement each other for the production of certain B vitamins (in particular biotin/B_7_ and riboflavin/B_2_) and essential amino acids [18–22,25–27]. These systems, involving distinct aphid species and strains of *S. symbiotica* acquired independently through evolution, are suitable to study the diversity of multi-symbiotic nutritional symbiosis and to lay the groundwork for studying the development and functioning of these associations.

In the present study, we sequenced and annotated the genomes of the symbionts *B. aphidicola* and *S. symbiotica* composing the di-symbiotic systems associated with two Chaitophorinae aphids: the cereal aphid *Sipha maydis* and the Norway maple aphid *Periphyllus lyropictus*. The prevalence of *S. symbiotica* reaches 100% in these two aphid species and our genomic analyses suggest that the symbiont is a recently acquired co-obligate partner. We found that the nutritional symbiont exhibits very contrasting infection patterns in the two aphid species: while *S. symbiotica* exhibits an invasive phenotype in *P. lyropictus*, the co-obligate symbiont is strictly compartmentalized into bacteriocytes in *S. maydis*. We suggest that these differences in the anatomical integration of the co-obligate symbiont can be explained by the way in which *S. symbiotica* and *B. aphidicola* complement each other metabolically in their system, but also by the set of virulence factors that each co-obligate symbiont has retained over its evolution with its respective host. This work provides insight into the metabolic and anatomical integration of co-obligate symbionts in Chaitophorinae aphids in the early stages of co-obligate symbiosis.

## Materials and methods

### Prevalence of *S. symbiotica* in *S. maydis* populations

The prevalence of a symbiont in insect populations is an indication of its symbiotic status, with a prevalence of 100% indicating that the symbiont is fixed in the host species concerned and is an obligate partner. To examine the prevalence of *S. symbiotica* in *S. maydis* populations, specimens were collected on three common cereals grown in Morocco (*Triticum turgidum, Triticum aestivum* and *Hordeum vulgare*) from the main cereal growing areas of this country in two sampling campaigns: one in April 2014 covering six regions and locations with 21 colonies sampled and one in April-May 2016 covering two regions with 76 colonies sampled (Figure S1; Table S1). The aphids collected consisted only of wingless parthenogenetic adult females that were stored in 95% ethanol at 4°C until use.

Prior to DNA extraction, insect samples were surface sterilized with 99% ethanol, 10% bleach and rinsed with sterile water. Genomic DNA was extracted using the DNeasy Blood & Tissue kit (Qiagen) following the manufacturer’s instructions. Each DNA extraction was performed on one individual per colony. DNA extractions were evaluated qualitatively and quantitatively using a Nanodrop spectrophotometer (Thermo Scientific) and stored at -20°C prior to PCR testing. Aphid samples were tested for the presence of *S. symbiotica* by amplifying a partial region of the 16S rRNA gene (0.4 kb region) using the specific primers 16SA1 and PASScmp [28]. PCR assays were performed as described previously [29]. The PCR products were stained with ethidium bromide and visualized on a 1% agarose gel. The amplicons were purified, sequenced, and then their identity was confirmed by BLAST search.

### Establishment of aphid clonal lines and rearing conditions

To perform the genomic approaches and other analyses, we established clonal lines for *S. maydis* and for *P. lyropictus*. For *S. maydis*, a colony sampled on *H. vulgare* in Midelt, (Morocco) in April 2016 was used to generate a clonal lineage from a single individual (clone Midelt). Aphids were reared on *Triticum aestivum* (bread wheat) under long day conditions (16h light, 8h dark) in a room maintained at a constant temperature of 20°C to ensure parthenogenic reproduction

The clonal line of *P. lyropictus* was established from one individual collected from *Acer platanoides* (Norway maple) in Louvain-la-Neuve in spring 2020 (clone LLN) [30]. Aphids were reared on young *A. platanoides* trees under the same conditions as the *S. maydis* clone Midelt.

### Genome sequencing and assemblies

For whole-genome sequencing, DNA samples enriched with bacteria from *S. maydis* and *P. lyropictus* clone lines maintained in the laboratory were prepared following a previously described protocol [20,31]. Two sequencing approaches were performed on the DNA extracts: MinION long-read sequencing (Oxford Nanopore approach) and Illumina short-read sequencing. Both approaches were used to generate a hybrid assembly and thus obtain the most complete and accurate genomes possible.

Prior to the MinION nanopore sequencing, the integrity of genomic DNA was evaluated using the 5200 Fragment Analyzer System (Agilent) with the Genomic DNA 50kb kit (Agilent). Libraries were then prepared from 400 ng of genomic DNA following the Rapid Barcoding Sequencing kit (SQK-RBK004) protocol (Oxford Nanopore Technologies). Briefly, the DNA molecules were cleaved by a transposase and barcoded tags were attached to the cleaved ends. The barcoded samples were then pooled and Rapid Sequencing Adapters were added to the tagged ends. The pooled libraries were sequenced into a FLO-MIN106 (R9.4.1) flow cell for a 48h run according to manufacturer’s instruction. After the run, fast5 files were basecalled on the MinIT using default settings in MinKNOWv21.02.5 and Guppy v4.3.4 and a Fast Basecalling. For the Illumina sequencing, libraries were prepared starting from 30 ng of genomic DNA using the Illumina DNA prep kit (Illumina, USA) following the manufacturer’s instruction. Briefly, genomic DNA was tagmented using Bead-Linked Transposomes which fragments and tags the DNA with adapter sequences. After a post tagmentation clean-up, tagmented DNA were amplified using a 12 cycle PCR program and IDT® for Illumina® DNA/RNA UD Indexes Set A to adds the index and adapter sequences. Libraries were equimolarly pooled and paired-end (2 × 300 bp) sequenced on an Illumina MiSeq platform. (Illumina).

Raw sequences generated by MinION nanopore sequencing were assembled using meta-flye v.2.9, an assembler supporting metagenomic data that was specifically developed for the assembly of third-generation long-read sequences [32]. The resulting draft assemblies were then polished with Illumina short reads by using multiple rounds of Pilon v1.24 [33]. Four genomes were thus assembled: those of the *B. aphidicola* and *S. symbiotica* symbionts composing the di-symbiotic system of *S. maydis* (hereafter BaSm and SsSm) and those of the *B. aphidicola* and *S. symbiotica* symbionts of *P. lyropictus* (hereafter BaPl and SsPl).

### Genome annotation and comparative analyses

The four assembled genomes were annotated with the Prokaryotic Genome Annotation Pipeline (PGAP) [34]. The genome sequences were deposited on the MicroScope platform [35,36] and their metabolic capabilities compared using MicroCyc, a collection of microbial Pathway/Genome Databases (PGDBs) based on the MetaCyc database and specifically dedicated to the analysis of microbial pathways [37]. Metabolic reconstructions generated by MicroCyc were manually verified and CDSs were considered as pseudogenes when their length was less than 80% of the one of their orthologs in reference bacterial genomes [38,39].

Virulomes of both *S. symbiotica* symbionts were analyzed with the virulence factor database (VFDB, http://www.mgc.ac.cn/VFs/) [40] integrated on the MicroScope platform using BLASTp (with a threshold of minimum 50% aa identity, 80% align. Coverage). Secretion systems were identified with MacSyFinder [41]. PHASTER was used to predict the intact prophage regions [42]. Assemblies and annotations are available on GenBank (BioProject accession numbers: PRJNA835797 for BaSm, PRJNA837269 for SsSm, PRJNA837360 for BaPl and PRJNA837363 for SsPl.

### Molecular phylogenetic analyses

To examine the evolutionary relationships of SsSm and SsPl with other *S. symbiotica* symbionts and other bacteria in the genus *Serratia*, a phylogenetic tree was built using a set of single-copy core concatenated protein sequences shared across all *S. symbiotica* strains for which complete genome sequences are publicly available. *S. symbiotica* and outgroup genomes used for phylogenetic analysis are listed in Table S2 in the supplemental material. Genomes were downloaded from the NCBI Assembly Database on June 2^nd^ 2022. All protein sequences of the different species and strains were analyzed with OrthoFinder (version 2.0) [43,44] to identify groups of orthologs. A set of 235 protein sequences encoded by single-copy genes present in each bacterial strain was then used for phylogeny estimation. The alignments of this set of protein sequences were concatenated using Phyutility [45]. By initially defining each protein as a separate partition, we performed a ModelFinder analysis [46] on the concatenated alignment with IQ-TREE (version 1.6.3) [47]; (options -m TESTMERGEONLY and -rcluster 10; set of models evaluated: mrbayes) to select the best partitioning of the alignment and the best model for each partition (-spp option, allowing each partition to have its own evolution rate). A maximal likelihood analysis was then conducted on the best partitioning and modelling scheme (-spp option) with IQ-TREE (options -bb 1000 and -alrt 1000 for 1000 replicated using ultrafast bootstrapping (UFBoot; [48,49]) and approximate likelihood ratio test (SH-aLRT; [50]), respectively.

### Examination of the tissue tropism of *S. symbiotica*

Tissue tropism of SsSm in *S. maydis* and of SsPl in *P. lyropictus* was examined using whole-mount fluorescence in situ hybridization (FISH) as described previously [51]. Young adult aphids were collected and preserved in acetone until fixation. For fixation, aphids were first placed in 70% ethanol where the legs were removed with micro-scissors, then transferred to Carnoy solution and left at room temperature overnight. After a thorough washing with 100% ethanol, the aphids were bleached in alcoholic 6% H_2_O_2_ solution and incubated at room temperature for two weeks. The bleached samples were successively washed with 100% ethanol and 70% ethanol, then hydrated with PBSTx containing 0.3% Triton X-100 and finally incubated with hybridization buffer (20 mM Tris-HCl [pH 8.0], 0.9 M NaCl, 0.01% SDS, 30% formamide) containing 100 nM of each probe and 0.5 μM SYTOX Green (Thermo Fisher Scientific) overnight. For *S. maydis*, the following oligonucleotide probes were used for in situ hybridization: Cy5-ApisP2a (5’-Cy5-CCTCTTTTGGGTAGATCC-3’) targeting 16S rRNA of *B. aphidicola* and Cy3-PASSisR (5’-Cy3-CCCGACTTTATCGCTGGC-3’) targeting 16S rRNA of *S. symbiotica* [52]. For *P. lyropictus*, the probe Cy5-PeriBuch (5’-Cy5-CCTTTTTTGGGCAGATTC-3’) was specifically used to target 16S rRNA of *B. aphidicola* [22]. After this incubation period, the samples were washed thoroughly with PBSTx, mounted in SlowFade antifade solution (Thermo Fisher Scientific), and observed under a Zeiss LSM 710 confocal microscope equipped with Airyscan detector.

## Results and discussion

### *S. symbiotica* is a fixed symbiont in *S. maydis*

Examination of the prevalence of *S. symbiotica* in *S. maydis* populations shows that all 97 colonies surveyed were positive for the symbiont (Table S1), indicating that, in this aphid species, *S. symbiotica* is a fixed symbiont involved in a stable relationship with its host. Indeed, unlike facultative symbionts that show irregular prevalence in host insect populations because they are not essential to host survival, are often associated with fitness costs, and tend to exhibit imperfect maternal vertical transmission [53–55], obligate symbionts are universally present in the host species because both partners have evolved an interdependent relationship [12]. The universal presence of *S. symbiotica* in *S. maydis* suggests that it is an obligate symbiont, as previously reported in other aphid species of the subfamilies Lachninae and Chaitophorinae [19–22,27]. A 100% prevalence of *S. symbiotica* in *P. lyropictus* populations has been reported previously [22,30], indicating that the symbiont is also fixed in this Chaitophorinae aphid.

### Genome sequencing of the *B. aphidicola*/*S. symbiotica* consortium

Sequencing of the *B. aphidicola*/*S. symbiotica* consortium genomes associated with each aphid species was performed using MinION sequencing (Oxford Nanopore approach) and Illumina MiSeq paired-end sequencing. For each technology, the total number of reads yielded, the average read length, and average genome coverage of each symbiont are summarized in Table S3. Using this hybrid assembly approach, highly contiguous de novo assemblies were obtained for each symbiont genome. The chromosomes of the *S. maydis*-associated *B. aphidicola* strain (BaSm) and the *P. lyropictus*-associated *B. aphidicola* strain (BaPl) could be circularized. The assembly approach also provided a circularized chromosome for the *S. symbiotica* co-obligate symbiont of *P. lyropictus* (SsPl), and identified an associated plasmid, namely pSsPl-LLN. The genome of the *S. symbiotica* strain associated with *S. maydis* (SsSm) consists of three contigs.

### General genomic features of the different symbionts

The general genomic features of the different symbionts targeted in this study are summarized in Table 1. Both *B. aphidicola* genomes are approximately 0.46 Mb in size, showing that the genomes of strains associated with *S. maydis* and *P. lyropictus* are much more eroded than *Buchnera* strains involved in mono-symbiotic nutritional systems (only *Buchnera* as nutritional symbiont), which are typically featured by a genome size of about 0.64 Mb [56]. These *Buchnera* symbionts with larger genomes are typically associated with aphids of the subfamily Aphidinae. In contrast, *B. aphidicola* symbionts with “small genomes” (i.e., around 4.5 Mb) are systematically found in aphids where this primary symbiont forms a metabolic consortium with a more recently acquired nutritional symbiont, typically in the Lachninae and Chaitophorinae aphids (Manzano-Marín and Latorre, 2014; Manzano-Marín et al., 2016, 2018; Manzano-Marín et al., 2020; Monnin et al., 2020). The small genome size of BaSm is thus further evidence suggesting that the cereal aphid *S. maydis*, similarly to the Norway maple aphid *P. lyropictus*, hosts a di-symbiotic system where *B. aphidicola* is metabolically complemented by a *S. symbiotica* co-obligate symbiont.

**Table 1.**
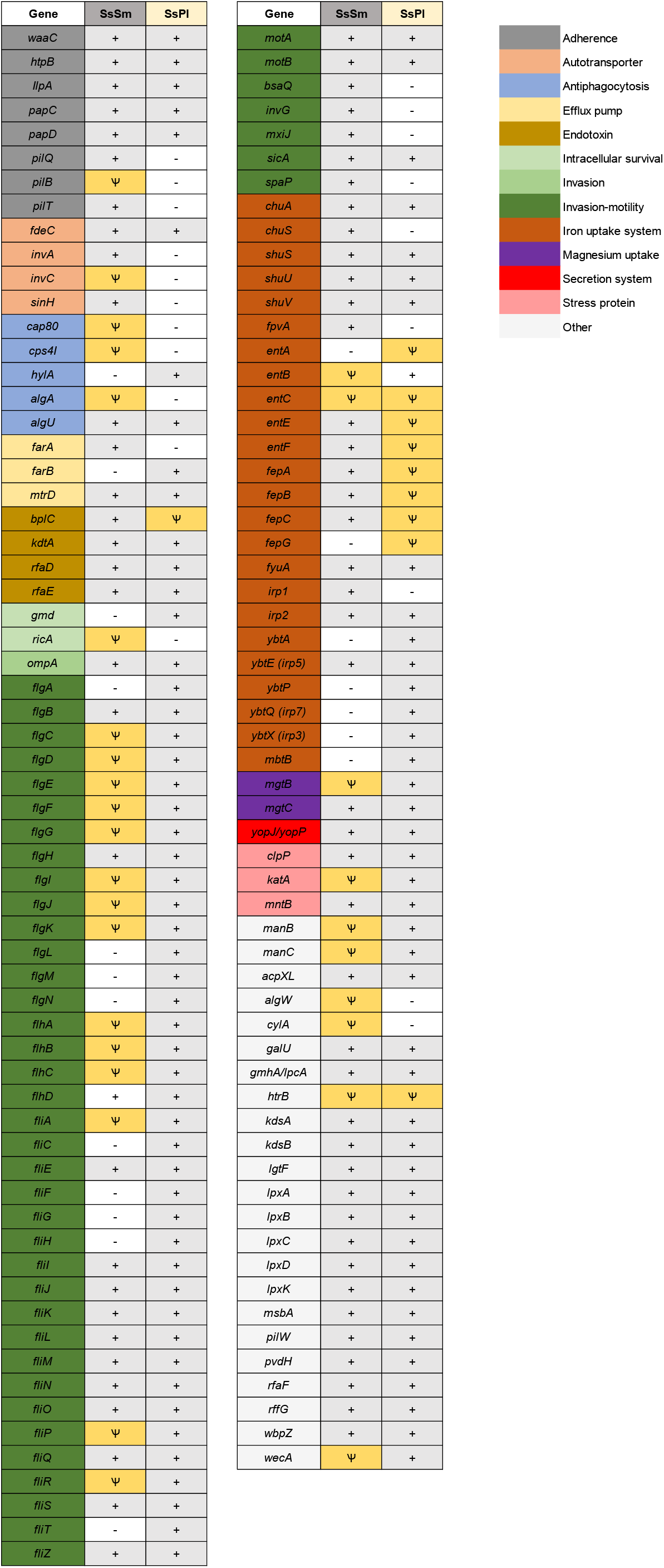
General genomic features of *B. aphidicola* and *S. symbiotica* symbionts associated with *S. maydis* and *P. lyropictus* respectively.

The genomes of the two *S. symbiotica* symbionts differ in size: whereas the SsSm genome consists of a 2.48 Mb genome carrying 1360 presumed intact CDSs, the SsPl genome consists of a 3.05 Mb, chromosome complemented by a large plasmid of 97.8 kb, carrying 2114 presumed intact CDSs. The genome size we report here for the *S. symbiotica* symbiont associated with *P. lyropictus* is larger than that reported by other authors in a previous study (2.58 Mb) that used another sequencing approach not compatible with chromosome circularization and plasmid identification [22]. This suggests that the version we report here provides a more accurate representation of the genome of the *S. symbiotica* co-obligate symbiont associated with *P. lyropictus*. Our data suggest that SsSm and SsPl were recently acquired by their host as they have genomes larger than 2.48 Mb, which is quite large for *S. symbiotica* co-obligate symbionts. For example, the *S. symbiotica* co-obligate symbiont associated with the conifer aphid *Cinara cedri* (Lachninae subfamily) has a genome size of 1.76 Mb and the genome of the strain associated with the giant willow aphid *Tuberolachnus salignus* (Lachninae subfamily) is 0.65 Mb in size [57]. The genomes of SsSm and SsPl are actually similar in size to those of the facultative *S. symbiotica* strains reported in *A. pisum* (SsApIS and SsApTucson) and the *S. symbiotica* co-obligate symbionts associated with *Cinara tujafilina* (SsCt) and *Cinara strobi* (SsCs) reported as recently acquired co-obligate symbionts [20,26,57]. The still large genome size of SsSm and SsPl thus indicates that they are still in the early stages of genomic reduction, especially SsPl which retains a larger genome. The hypothesis of recently integrated *S. symbiotica* symbionts as obligate nutritional partners is also supported by the presence in both genomes of a high proportion of pseudogenes, 1258 pseudogenes/2617 total CDSs (48%) in the SsSm genome and 1172 pseudogenes/3286 total CDSs (36%) in the SsPl genome. The presence of a high proportion of pseudogenes is an indicator of massive loss of functional genes and is another genomic feature typical of the early stages of genome reduction and evolutionary transition towards a host-dependent lifestyle [58]. The proportion of pseudogenes is much higher in the SsSm genome than in the SsPl genome, indicating that SsSm is probably more advanced in the evolutionary process of genomic degeneration than SsPl. This hypothesis is also supported by the smaller genome size of SsSm as well as the lower proportion of intact phage regions compared to SsPl (Table 1; Table S4).

Finally, it is interesting to note that, at 3.15 Mb (including chromosome and plasmid), the SsPl genome is the largest *S. symbiotica* genomes described so far for an aphid co-obligate symbiont. It is also larger than any genome of facultative *S. symbiotica* symbiont reported to date and approaches the genome size of cultivable pathogenic strains that have been isolated from aphids of the genus *Aphis* [59–61] and associated with a propensity to rapidly invade the aphid digestive tract [29,57,61–63]. This support the hypothesis that, in the evolution of bacterial mutualism, SsPl is at a very early stage of its establishment as a vertically transmitted obligate symbiont [30].

### Phylogenetic positioning of SsSm and SsPl and tissue tropism

We examined the evolutionary relationships of SsSm and SsPl with all *S. symbiotica* strains for which complete genomic sequences are publicly available. Our phylogenetic analysis is based on 235 shared orthologous genes and is rooted with outgroups that included other members of the genus *Serratia* and *Yersinia pestis* as a more distant Enterobacterales species. Our phylogenomic tree shows that the *S. symbiotica* strains fall into two clades (Figure 1), as previously established [61,64]. Both SsSm (Midelt) and SsPl (LLN) are part of clade A, which is composed of strains that act as pathogens or mutualists in aphids of the subfamily Aphidinae, but also includes co-obligate strains associated with Chaitophorinae aphids and the co-obligate strain associated with *Cinara tujafilina* (subfamily Lachninae). Clade B includes only co-obligate strains associated with aphid species of the subfamily Lachninae.

**Figure 1.**
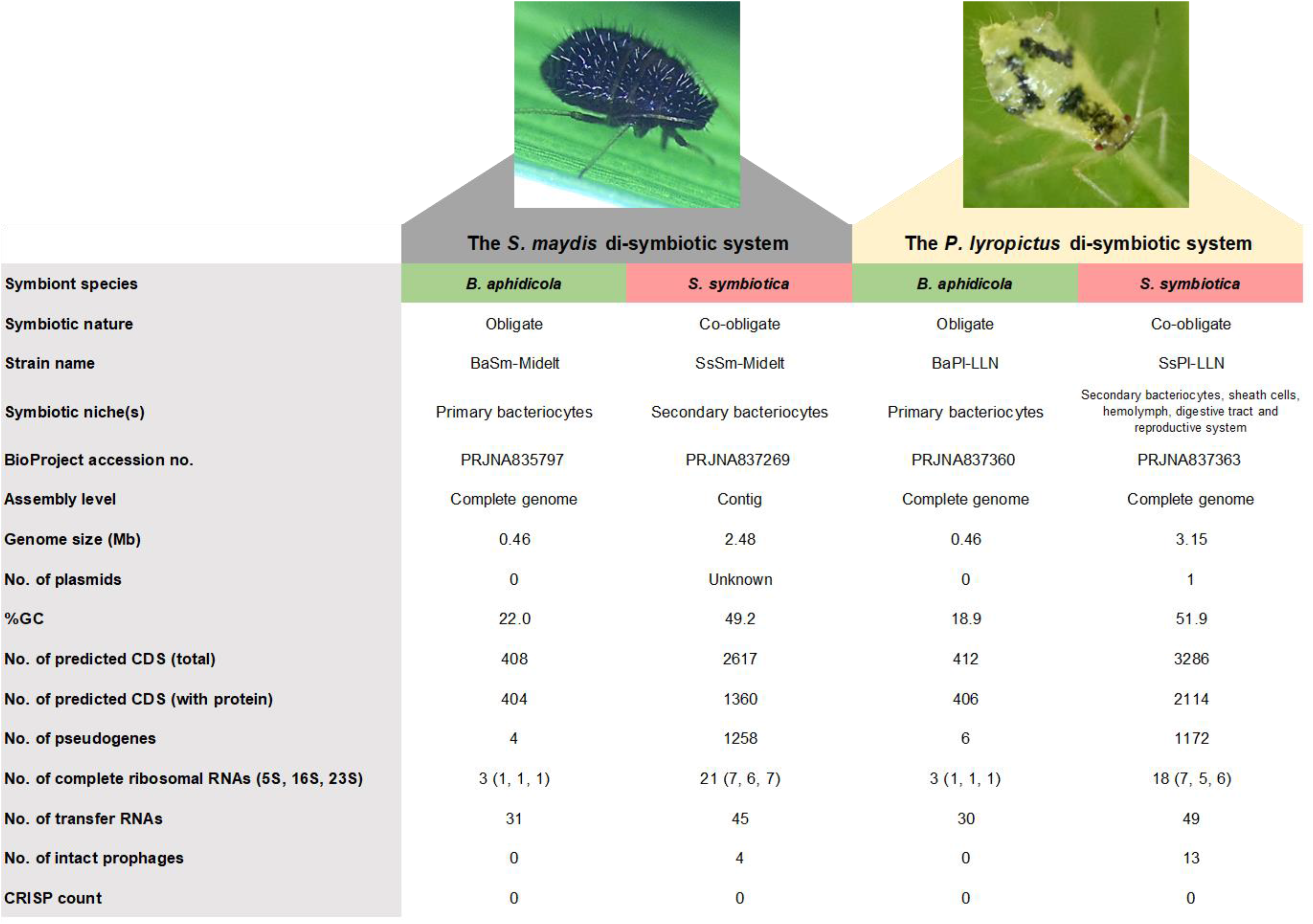
Maximum likelihood phylogeny of *S. symbiotica* based on concatenation of 235 single-copy orthologs shared across *S. symbiotica*, other *Serratia* species and *Yersinia pestis*. Branch support (SH-aLRT and ultrafast bootstrap values) was >90% for all but one node*: *S. symbiotica* Af-2.3, *S. symbiotica* Af-24.1, *S. symbiotica* Apa-8A1 and *S. symbiotica* Ct. The *S. symbiotica* strains composing clade A are highlighted in blue while the strains composing clade B are highlighted in green. The outgroup is highlighted in gray. Strains SsSm-Midelt and SsPl-LLN are denoted in bold red. Genomes used for the phylogeny and relevant references for *S. symbiotica* genome features are listed in Table S2 in the supplemental material.

Regarding the tissue tropism of the *S. symbiotica* co-obligate symbionts in the two aphid species, it is marked by very contrasting localization patterns. SsPl exhibits a dual mode of life: intracellular by being housed in large syncytial secondary bacteriocytes embedded between the primary bacteriocytes containing *B. aphidicola* in a well-organized compartmentalization pattern, and extracellular by invading a wide variety of host tissues, including the digestive tract, hemolymph and oviduct (Figure 2). The tissue tropism and infection dynamics of SsPl during *P. lyropictus* development have been studied in detail previously [30]. In contrast, SsSm appears to be only compartmentalized within the large secondary bacteriocytes embedded between the primary bacteriocytes hosting *B. aphidicola* and SsSm is not found in hemolymph or other tissues (Figure 2). Compartmentalization within bacteriocytes is a typical feature of obligate nutritional symbionts [65,66] and a key adaptation that underpins the partnership between insects and these bacteria. Indeed, these specialized host cells mediate metabolic exchanges between the host and symbiotic bacteria and allow the host to control populations of its symbionts according to its nutritional needs [2,4,67]. The *S. symbiotica* co-obligate symbiont associated with *P. lyropictus*, exhibits invasive traits and escape strict compartmentalization into bacteriocytes despite its integration into a cooperative lifestyle with its host and the ancestral symbiont *B. aphidicola* [30]. It has been suggested that the invasive phenotype of SsPl could be explained by the very recent acquisition of the symbiont by its host, which is not yet integrated into a fully stabilized relationship. The evolutionary history of SsSm with its host is likely more ancient, resulting in further anatomical integration, i.e. strict compartmentalization of the co-obligate symbiont into *S. maydis* bacteriocytes. SsSm likely lost the ability to adopt an extracellular lifestyle, probably as a result of the loss of the genes required for it on the road to reductive evolution and which constrains it to an intracellular lifestyle. Such an anatomical configuration is expected to support the metabolic exchanges between *B. aphidicola* and *S. symbiotica*. Given the contrasting tissue tropism of *S. symbiotica* between *P. lyropictus* and *S. maydis* (invasive or strictly compartmentalized), it is likely that host-specific mechanisms are used to control populations of the nutritional symbiont. In particular, host tolerance to the highly invasive nature of SsPL requires further study.

**Figure 2.**
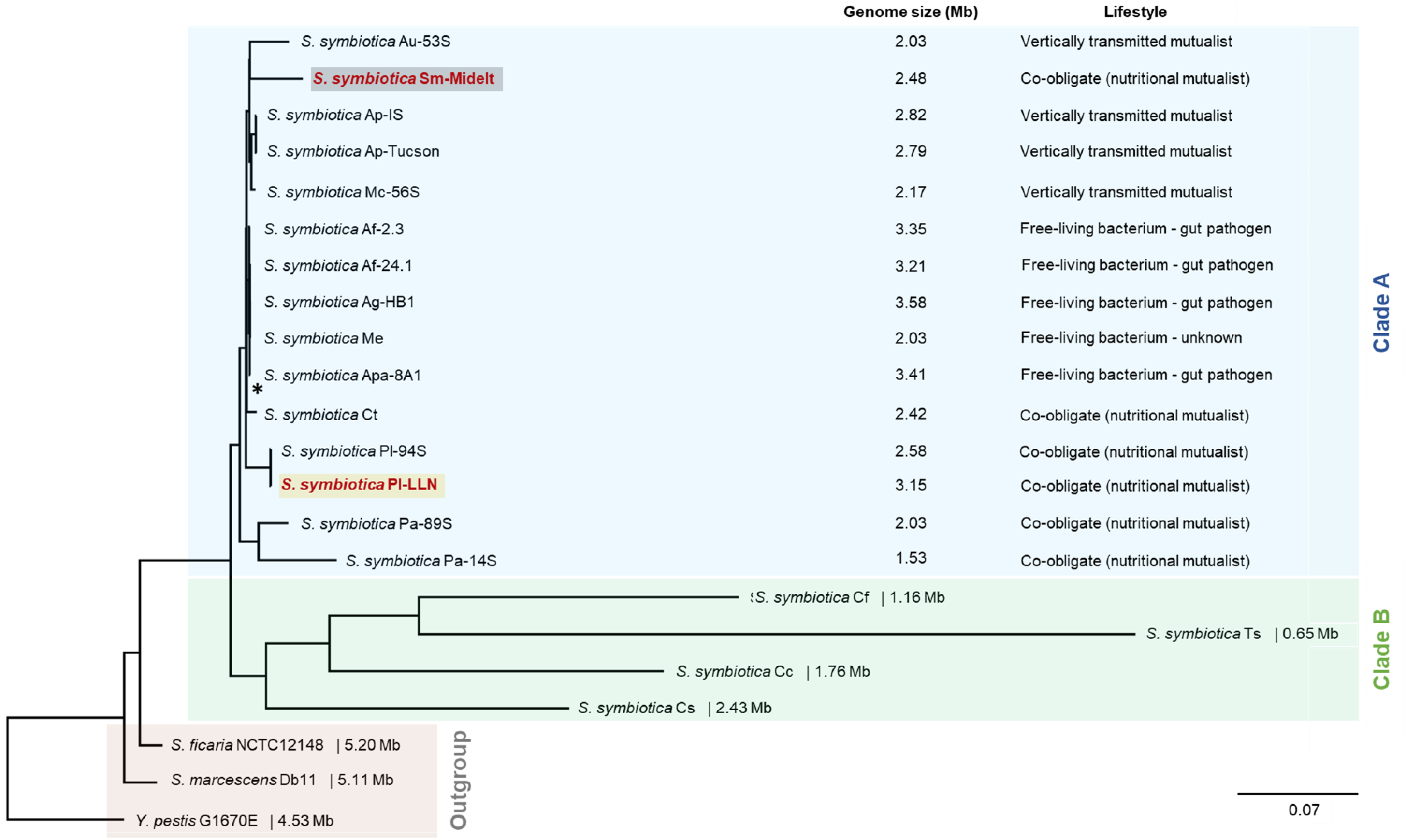
Tissue tropism of the *S. symbiotica* co-obligate symbiont associated with *P. lyropictus* and with *S. maydis*. Green, red, and blue signals indicate *Buchnera* cells, *Serratia* cells, and host insect nuclei, respectively. (A) In *P. lyropictus*, the *S. symbiotica* co-obligate symbiont resides extracellularly in the gut and intracellularly in secondary bacteriocytes (Sba) embedded between primary bacteriocytes (Pba) harboring the primary symbiont *Buchnera*. These clusters are distributed along the aphid’s abdomen. *S. symbiotica* can also be found in the gut of embryos (see embryo outlined in yellow frame) (B) *S. symbiotica* massively infecting the gut (here a portion of the midgut). (C) One of the bacteriocyte clusters forming the bacteriome composed of syncytial secondary bacteriocytes hosting *S. symbiotica*, embedded between the uninucleated primary bacteriocytes hosting *Buchnera*. Sheath cells that also house the co-obligate symbiont sparsely cover the periphery of the bacteriocytes (arrowheads). (D) *S. symbiotica* massively infect the periphery of the embryos in the ovarioles. The tissue tropism and infection dynamics of *S. symbiotica* in *P. lyropictus* has been thoroughly mapped previously [30]. (E) In *S. maydis*, the bacteriome has a horseshoe shape and consists of secondary bacteriocytes hosting *S. symbiotica* embedded between the primary bacteriocytes hosting *Buchnera*. (F) In *S. maydis*, the *S. symbiotica* co-obligate symbiont is strictly compartmentalized into secondary bacteriocytes. Unlike SsPl, SsSm does not evolve in the digestive tract of its host, nor in the hemolymph and does not appear to reside in the sheath cells. (G) Close-up view of one of the lobes of the bacteriome showing *S. symbiotica* only present in the secondary syncytial bacteriocytes.

### In the two Chaitophorinae aphids, *S. symbiotica* and *B. aphidicola* complement each other metabolically but in distinct layouts

In many Lachninae and Chaitophorinae aphids, the ancestral primary symbiont *B. aphidicola* has become unable to synthesize some essential nutrients on its own due to drastic genome degradation, and the synthesis of these nutrients is supported by the completion of *Buchnera*’s missing pathways with those of a more recently acquired co-obligate symbiont, the whole forming a metabolic unit [18,22,26,27]. How the co-obligate symbiont metabolically complements *B. aphidicola* is usually inferred by genomic analyses. In Lachninae and Chaitophorinae aphids, the co-obligate symbiont is typically required for the biosynthesis of the vitamins riboflavin (B_2_) and biotin (B_7_) [20–22,25,26]. Analysis of the di-symbiotic systems of *S. maydis* and *P. lyropictus* shows that in both cases *S. symbiotica* likely takes charge of riboflavin biosynthesis (Figure 3). Indeed, unlike what has been observed in mono-symbiotic aphids [21], *B. aphidicola* in the studied Chaitophorinae aphids did not conserve any of the genes for synthesis of this vitamin. In contrast, all these genes are present in the genome of co-obligate symbionts, except for the gene encoding the 5-amino-6-(5-phospho-d-ribitylamino)uracil phosphatase enzyme in SsSm. *yigB* and the alternative gene *ybjI* are absent from the genome of SsSm and SsPl. *yigL*, reported as another alternative for this phosphatase activity [68] is present and intact in the SsPl genome, but pseudogenized in the SsSm genome (split into four pieces). The genes known to encode this enzymatic activity can be missing from the genome of *Buchnera* strains involved in mono-symbiotic systems (e.g., the APS strain) [69]. However, it has been postulated that this phosphatase activity may be performed by alternative enzymes whose identity is still unclear [68,70]. Regarding biotin biosynthesis, our results show contrasting biosynthetic capacities between the two co-obligate symbionts SsSm and SsPl. SsSm appears to complement *B. aphidicola* for the biosynthesis of this vitamin from 7-keto-8-aminopelargonate. Indeed, the genes *bioA, bioB* and *bioD* are intact in the genome of the co-obligate symbiont associated with *S. maydis*, as is the case in the co-obligate symbionts associated with Lachninae aphids [21]. In contrast, *bioA* and *bioD* are pseudogenized in SsPl. The case of the di-symbiotic system associated with *P. lyropictus* for biotin biosynthesis thus differs from what has been previously reported for other Lachninae and Chaitophorinae aphids in which the genes *bioA, bioB*, and *bioD* of the co-obligate symbiont are intact [18,21,22]. One hypothesis is that the phloem sap of the Norway maple tree on which *P. lyropictus* specifically feeds contains this vitamin in sufficient quantities for the aphid to dispense with the biosynthetic capabilities of its bacterial partners.

**Figure 3.**
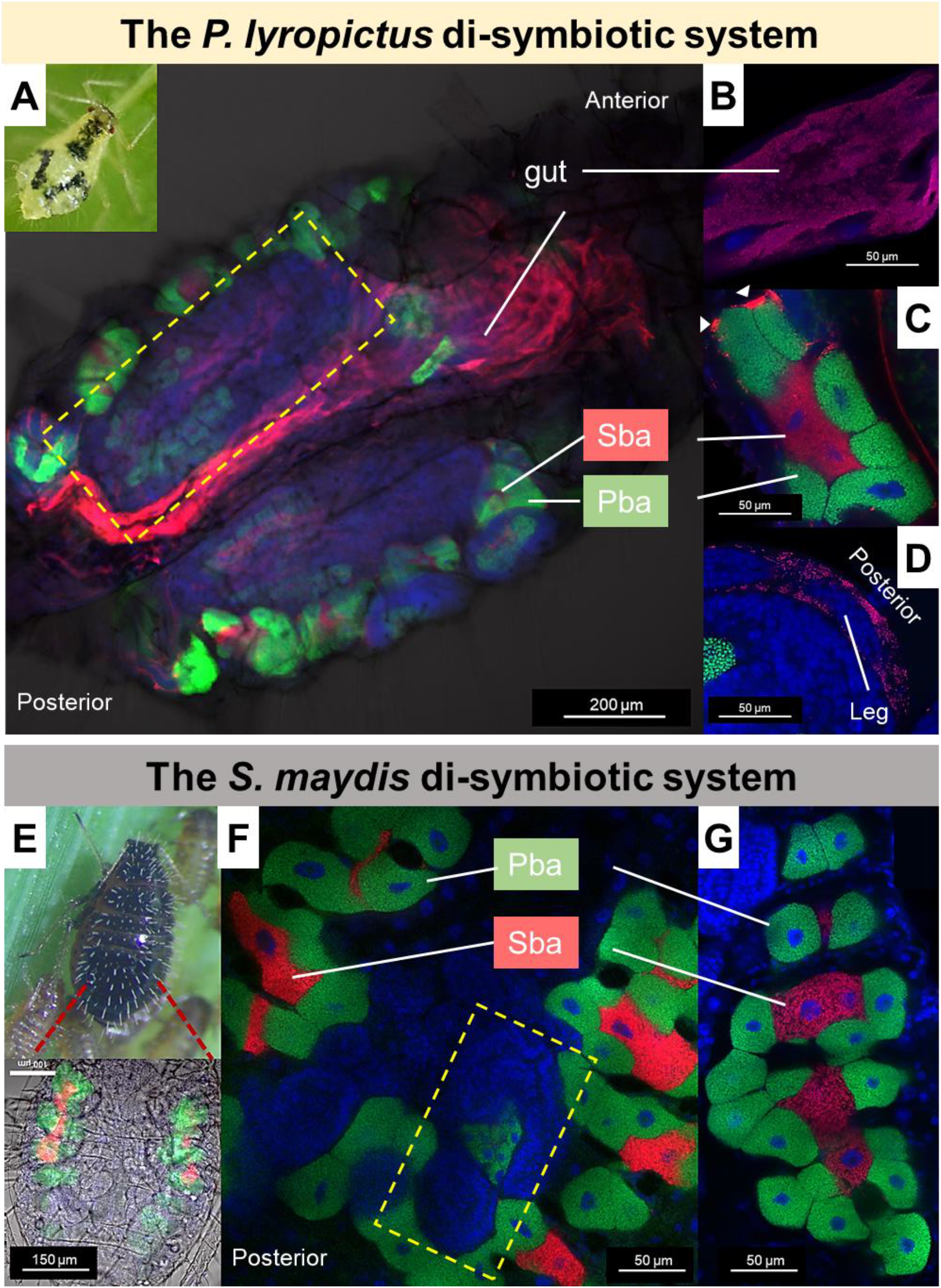
B_2_ and B_7_ vitamins biosynthetic metabolic capabilities of the di-symbiotic system *B. aphidicola* (Ba)-*S. symbiotica* (Ss) in *S. maydis* (Sm) and *P. lyropictus* (Pl). The *Buchnera* APS strain forming the mono-symbiotic system (*Buchnera*-only) associated with the pea aphid *A. pisum* was used as a comparison. On the left are the names of the genes encoding the enzymes involved in the biosynthetic pathway. Each box is associated with the gene encoded by the genome of a symbiont. The following code is used to characterize the state of each gene: the blue color means that the gene (or an alternative gene that can perform the same enzymatic function) is present and not pseudogenized; the gray color indicates that the gene is missing; the yellow color with the psi (Ѱ) symbol means that the gene is pseudogenized (including its alternative(s)).

Regarding essential amino acid (EAA) biosynthesis (Figure 4), the two di-symbiotic systems also show contrasting patterns. A notable finding is that in *S. maydis*, the co-obligate symbiont SsSm did not retain any intact genes for the synthesis of essential amino acids, except for lysine biosynthesis. In contrast, the *B. aphidicola* strain associated with this aphid, despite its highly eroded genome, fully supports the biosynthesis of several EAAs, including histidine and tryptophan. In *P. lyropictus*, histidine biosynthesis is performed by SsPl and the tryptophan pathway is split between *B. aphidicola* and the co-obligate symbiont in a similar fashion to what has been reported in the Lachninae aphid *Cinara cedri* [71]. Overall, our results indicate that SsSm has no complementary role to *B. aphidicola* in EAAs synthesis and has evolved a dependence on the primary symbiont for the acquisition of most EAAs. The opposite situation occurs in *P. lyropictus*. Indeed, fewer genes involved in EAA biosynthesis are missing or pseudogenized in the SsPl genome compared to that of SsSm. The *S. symbiotica* co-obligate symbiont of *P. lyropictus* has, for example, all the genes required for arginine, histidine, lysine, methionine and threonine biosynthesis and retains many redundant genes with BaPl. This broader capacity of SsPl for EAAs biosynthesis supports the hypothesis that this *S. symbiotica* symbiont was more recently acquired by its host compared to SsSm in *S. maydis*. In this scenario, relaxed selection on these genes may not yet have led to their pseudogenization or elimination in the context of reductive evolution.

**Figure 4.**
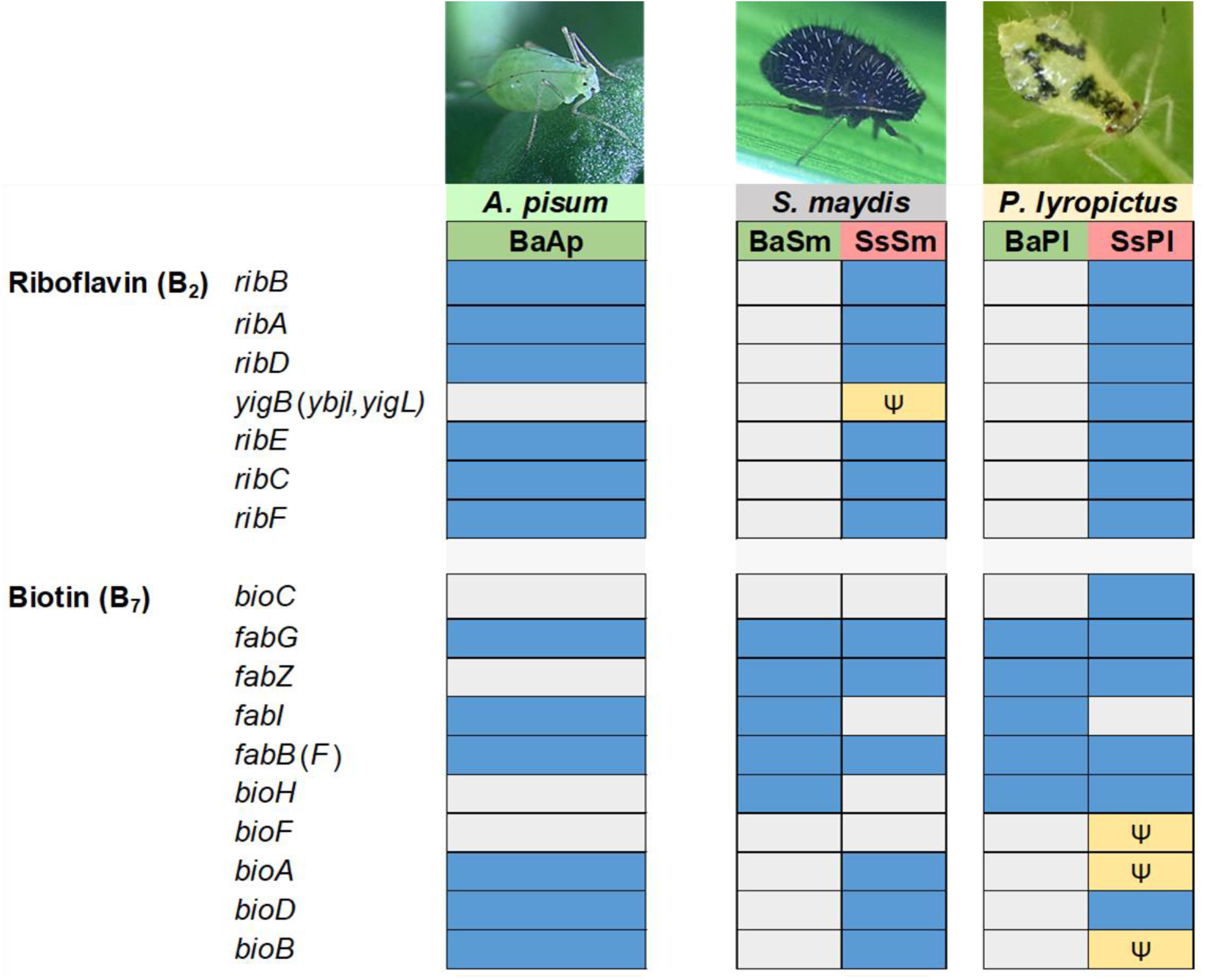
Essential amino acid biosynthetic metabolic capabilities of the di-symbiotic system *B. aphidicola* (Ba)-*S. symbiotica* (Ss) in *S. maydis* (Sm) and *P. lyropictus* (Pl). The *Buchnera* APS strain forming the mono-symbiotic system (*Buchnera*-only) associated with the pea aphid *A. pisum* was used as a comparison. On the left are the names of the genes encoding the enzymes involved in the biosynthetic pathway. Each box is associated with the gene encoded by the genome of a symbiont. The following code is used to characterize the state of each gene: the blue color means that the gene (or an alternative gene that can perform the same enzymatic function) is present and not pseudogenized; the gray color indicates that the gene is missing; the yellow color with the psi (Ѱ) symbol means that the gene is pseudogenized (including its alternative(s)).

In light of all these results, both SsSm and SsPl undertake riboflavin biosynthesis, which *B. aphidicola* is no longer able to perform. However, the way in which *B. aphidicola* and *S. symbiotica* complement each other metabolically for EAA biosynthesis differs greatly between the two di-symbiotic systems. One reason for this is that the EAA biosynthetic capabilities of the ancestral obligate symbiont *B. aphidicola* differ greatly between the two strains, in particular for histidine and tryptophan biosynthesis, whose pathways are complete in BaSm but incomplete in BaPl. This implies that, during the transition from facultative to co-obligate symbiont, the way in which relaxed selection on metabolism-related genes of the co-obligate symbiont is exerted depends on the genomic background of the primary symbiont and, in some cases, also that of the co-obligate symbiont it replaces. Added to this are the metabolic capabilities of the aphid host, which may differ between species, but which must play a role in the way di-symbiotic nutritional systems are shaped during evolution. This aspect has received little attention in studies of metabolic complementarity between symbionts, in part because it requires laborious sequencing and annotation of the host insect genome [72], a step that is however essential to establish the most accurate picture of the role of each partner in the functioning of a three-partner symbiotic system. Finally, another factor that future studies should consider is the nutrient content of the host plant phloem sap, that is expected to shape the evolution of specific biosynthetic pathways in nascent nutritional symbionts. Indeed, the nutrient content of phloem sap can differ greatly between plant species [73,74]. *S. maydis* and *P. lyropictus* are Chaitophorinae aphids that feed on contrasted host plants: while *S. maydis* feeds on the phloem sap of Poaceae (herbaceous plants), *P. lyropictus* feeds specifically on the phloem sap of the Norway maple *A. platanoides* (a woody plant). It is likely that the evolution of the di-symbiotic system associated with each of these aphid species is fashioned by the presumably very different nutritional content of the phloem sap of their respective host plant.

### The secretion systems and the array of virulence factors encoded by *S. symbiotica* genomes

The tissue tropisms of SsSm and SsPl suggest that the two co-obligate symbionts, although both performing a nutritional function, exhibit contrasting lifestyles within the host. Whereas SsSm is solely compartmentalized into secondary bacteriocytes, SsPl is capable of infecting a wide diversity of host tissues (Figure 2; [30]). These differences in invasion patterns should reflect the infection mechanisms that the two symbionts are able to express despite the ongoing reduction of their genome. The ability of bacteria to invade host tissues and cells depends in particular on the secretion systems they can express [75]. These protein complexes on bacterial cell membranes allow the translocation of effector proteins that can modulate the host environment and facilitate its invasion, for example by enhancing attachment to eukaryotic cells, plundering resources in an environmental niche, or intoxicating target cells and disrupting their functions. Our genome analyses reveal that SsSm and SsPl are associated with a low diversity of secretory systems (Table S5). The two *S. symbiotica* co-obligate symbionts retained only the ability to synthesize type V secretion systems (T5SS). These results are in agreement with those of a previous study showing that both facultative and co-obligate mutualistic strains of *S. symbiotica* do not retain complete secretion systems other than T5SS, in contrast to pathogenic strains that tend to retain key elements of T3SSs [57]. All *S. symbiotica* strains whose genomes have been sequenced so far, including those with extremely small genomes, are virtually capable of expressing T5SSs [57], raising the question of their possible role in the symbiosis with the insect host. T5SSs include diverse forms of virulence factor-secreting autotransporters involved in cell-to-cell adhesion and biofilm formation [76,77]. They may be involved in various functions associated with the specific lifestyle of the bacteria, including protease activity, intracellular mobility and acquisition of nutrient acquisition in limited environments. SsSm and SsPl appear to have retained the ability to synthesize a FdeC-like autotransporter adhesin (Figure 5; Tables S6). This T5SS has been shown to contribute to virulence of uropathogenic *Escherichia coli* (UPEC) strains by mediating their adhesion to mammalian cells [78], but the exact role of this autotransporter remains unknown outside this group of bacteria. Unlike SsPl, SsSm appears to have retained the genes encoding the autotransporters InvA (involved in the virulence of *Yersinia* bacteria [79]) and SinH (involved in the virulence of UPEC strains [80]). Currently, multiple functions of type V autotransporters have been identified recently in the context of eukaryotic host-pathogen interactions [76,81]. However, their potential roles in the context of bacterial mutualism in insects remain unknown.

**Figure 5.**
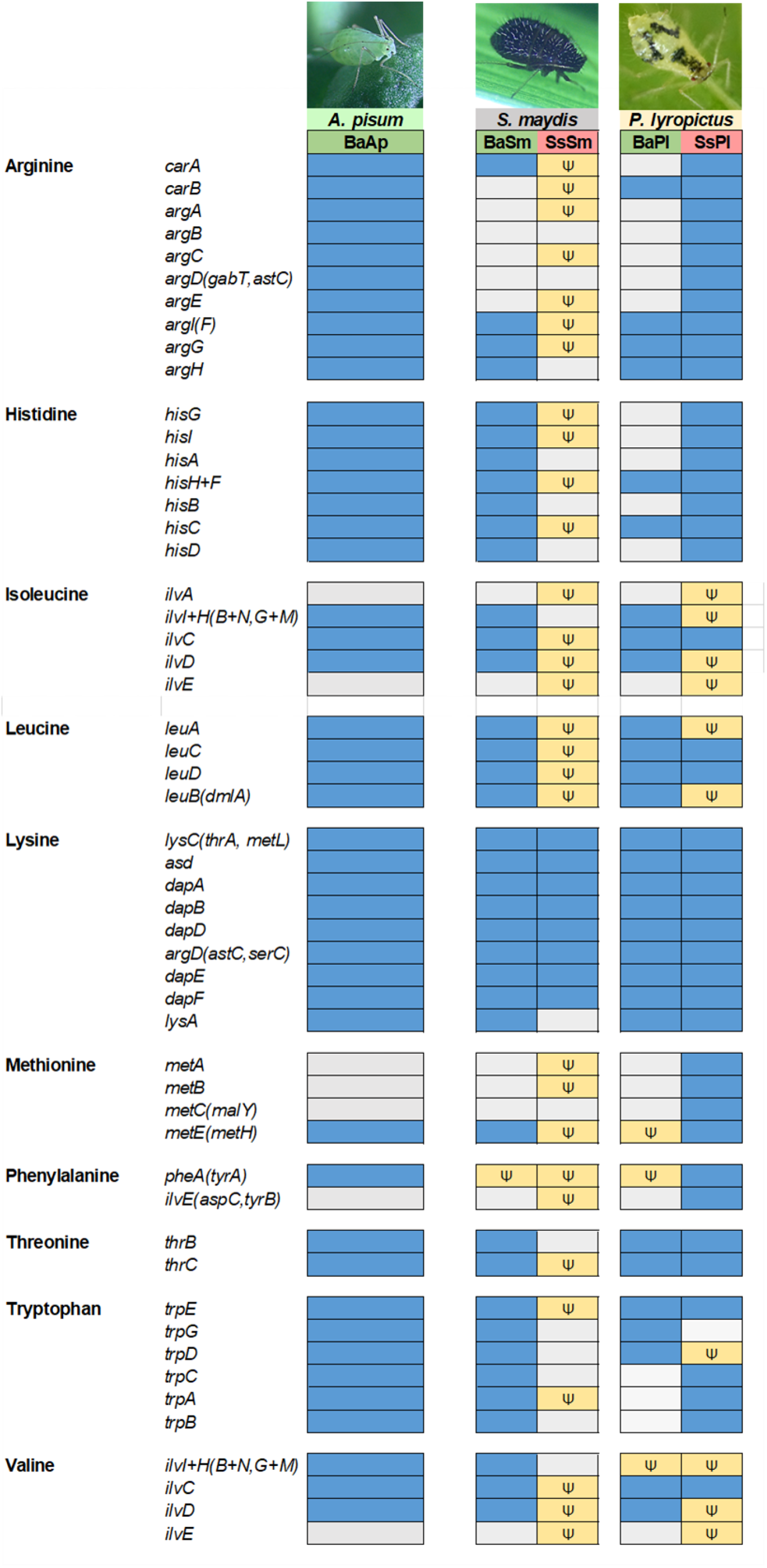
Presence/absence map of *S. symbiotica* virulence genes. Presence of a virulence gene is labeled in gray (+) and absence in white (-). The yellow color (Ѱ) means that the gene is pseudogenized. Note that the absence of a gene may be threshold dependent (minimum 50% aa identity, 80% align. Coverage).

The success of host colonization by invasive bacteria, whether in a host-pathogen relationship or mutualistic symbiosis, is largely determined by the ability of microorganisms to interact with hosts through the expression of virulence factors [82,83]. The content of virulence factors has been little studied in co-obligate symbionts. Yet, these nutritional symbionts, acquired more recently and exhibiting more moderate genome reduction compared to the ancient primary symbionts, are expected to have retained a greater diversity of these factors, which may allow them to colonize a greater diversity of host tissues. Examining virulence genes in the genome of co-obligate symbionts is therefore crucial to gain insight into the mechanisms underlying the early stages of insect endosymbiogenesis. In this context, our analyses show that SsSm and SsPl genomes differ greatly in their virulence factor content (Figure 5). Both genomes contain an equivalent number of virulence factor-encoding genes, but more genes are pseudogenized in SsSm than in SsPl (Figure 5; Table S6). This again suggests that SsSm is more advanced in reductive genome evolution compared to SsPl. The most striking example concerns the ability of both *S. symbiotica* strains to biosynthesize a complete flagellum, a macromolecular system that enables invasive bacteria to move through body fluids, attach the organ surfaces, and then promotes efficient colonization of host tissues [84]. The SsSm genome contains many genes encoding a flagellum, but those involved in hook and filament biosynthesis are either missing (including *fliC* and *flgL*) or pseudogenized (including *flgK, flgE, flgC, flgF* and *flgG*) (Table S6), suggesting that this structure has not retained any motility properties. Moreover, the remaining intact genes (including *flgB, flgH, fliE, fliI, fliM, fliN*) that correspond to the basal body of the flagellar structure, are also not sufficient for its function as an injectisome, a macromolecular protein complex whose structure is an evolutionary homologue of the bacterial flagellum and which is used by many bacteria, including bacterial endosymbionts, to deliver secreted effector proteins to a eukaryotic host [85,86]. The remnants of the flagellar apparatus associated with SsSm hardly allow for the biosynthesis of a reduced flagellum as found in some strains of *B. aphidicola* [85]. Motility pathways are among the first to be altered upon transition from a free-living to a host-dependent lifestyle [87]. Given that the SsSm genome is already highly reduced and pseudogenized, it is not surprising that this co-obligate symbiont has lost the capacity for motility.

In contrast, the SsPl genome contains all the genes theoretically required for biosynthesis of a complete flagellum (Figure 5; Table S6), suggesting that this *S. symbiotica* co-obligate symbiont is endowed with motility and chemotaxis potential. This could explain the ability of this *S. symbiotica* co-obligate symbiont to exhibit a highly invasive phenotype [30]. The case of SsPl, capable to adopt both an intracellular and an extracellular lifestyle, is singular for a nutritional symbiont, but not unique. Indeed, there are a few cases of bacteriocyte-associated nutritional symbionts that, despite the significant reduction of their genome and their specific nutritional function, retain a complete flagellar system [88,89]. *Wigglesworthia*, the obligate symbiont of tsetse flies, can adopt both an intracellular lifestyle in bacteriocytes and an extracellular phase in milk gland secretions during which it expresses a functional flagellum despite a severely reduced genome [88,90]. The case of SsPl is not as extreme as that of *Wigglesworthia*, but it illustrates the existence of co-obligate strains of *S. symbiotica* that, despite their integration in a cooperative system with the host and the primary symbiont *B. aphidicola*, resemble the previously described pathogenic strains in that they retain a complete flagellar system and the ability to colonize the host’s digestive tract [57]. One hypothesis is that SsPl express a functional flagellum depending on the tissue it colonizes and its lifestyle (i.e. intracellular or extracellular) as is the case for the symbiont *Wigglesworthia* in the tsetse fly.

Iron uptake by siderophores are well known to promote host colonization by pathogenic bacteria, but also by insect symbionts such as *S. glossinidius*, which uses this system to invade the tsetse fly [91]. Our analyses show that both *S. symbiotica* genomes contain many genes involved in siderophore biosynthesis. They both contain genes involved in the biosynthesis of enterobactin, a catechol-containing siderophore released by bacteria to bind ferric ions required for metabolic pathways [92]. However, in both SsSm and SsPl, most of the genes related to enterobactin biosynthesis (*entA, entB, entC, entE* and *entF*) are missing or pseudogenized, suggesting a relaxation of selection on these genes and that the production of enterobactin-like siderophores became dispensable during the evolution of both symbionts or was a limiting factor in the establishment of a persistent relationship between *S. symbiotica* and the host (Figure 5). Our analyses show that both symbionts retain intact genes for yersiniabactin (Ybt) biosynthesis, for which many identified genes are intact (e.g. *irp1, irp2, fyuA* and *ybtA*), especially in SsPl genome. However, despite many intact genes in the SsPl genome for the *ybt* locus, close examination shows that the locus is incomplete with some genes missing, including *ybtS, ybtu, ybtT, ybtD* and *psn*. Thus, it appears that SsPl has become unable to synthesize yersinibactin.

Finally, our analyses detected few toxin-related genes in the genomes of both co-obligate symbionts. SsSm and SsPl appear, however, to be able to synthesize effectors of the YopJ family, which are well known to promote invasion of pathogenic bacteria, but whose potential roles in the establishment of the mutualistic symbiosis remain unknown [93].

Based on the analysis of the virulence factors of these two co-obligate symbionts, it can be concluded that SsSm and SsPl exhibit different sets of virulence factors that reflect their contrasting lifestyles. The most notable finding is that SsPl is potentially capable of synthesizing a complete flagellum that would endow it with motility, an ability that could explain the remarkable invasive phenotype of this co-obligate symbiont. However, both co-obligate symbionts tend to exhibit low virulence factor diversity compared to cultivable *S. symbiotica* and especially compared to entomopathogenic bacteria of the genus *Serratia* such as *Serratia marcescens* [57]. Our results suggest that SsSm and SsPl are derived from originally pathogenic bacteria capable of synthesizing enterobactin and yersiniabactin-like siderophores, an ability they seem to have lost. Loss of virulence factors is conducive to the establishment of a long-lasting, mutualistic lifestyle with the host, which underlies strong selection for attenuated virulence [7].

## Conclusion and perspectives

In conclusion, the *S. symbiotica* co-obligate symbionts associated with two Chaitophorinae species, *S. maydis* and *P. lyropictus*, provide a contrasting picture of recent co-obligate nutritional endosymbiosis in aphids. The di-symbiotic systems associated with these aphids show contrasting facets regarding genome evolution and the level of anatomical integration of the co-obligate symbionts into their respective hosts. The *S. symbiotica* co-obligate symbiont associated with *P. lyropictus* still has a large genome, comparable in size to that reported for pathogenic cultivable strains. Its dual intracellular and extracellular lifestyle indicate that host dependence on nutritional symbionts may evolve prior to compartmentalization in the host cells specialized for metabolic exchange and symbiont control (i.e., bacteriocytes) [30]. Its ability to infect a wide variety of host tissues is probably due to its ability to express a complete flagellum and thus display an ability to move in body fluids and on organ surface, a well-known invasive potential typically expressed by pathogenic bacteria. How such a nutritional symbiont, both compartmentalized into bacteriocytes and exhibiting such an invasive phenotype, is controlled by its host is a puzzle to be solved in future studies. The *S. symbiotica* co-obligate symbiont associated with *S. maydis* exhibits another side of co-obligate nutritional endosymbiosis, that of further anatomical integration into the host (i.e., strict compartmentalization into bacteriocytes), indicative of a presumably older association with its host compared to SsPl, and now stabilized. In recent years, a growing number of studies have shown that many insect species harbor nutritional multi-symbiotic systems. However, how they develop and function remains largely unknown. Studying the diversity of these associations is crucial to tackle these aspects and address key questions in the evolutionary developmental biology of endosymbiosis including: How do metabolically complementary symbionts, forming a metabolic unit, coordinate with each other and with the host? How does the host control these nutritional symbionts, sometimes associated with an invasive phenotype? How do the different types of bacteriocytes (primary versus secondary) develop and interact with each other? The diversity of the di-symbiotic systems associated with Chaitophorinae aphids provides an ideal playing field to address these questions and better appreciate the sophistication of bacterial mutualism in insects.

## Acknowledgments

The authors thank Karen Gaget for her help with the aphid pictures and the whole BF2I laboratory for the stimulating discussions about the topic. This study was financially supported by FNRS grant no. 1B374.21. This paper is publication BRC 393 of the Biodiversity Research Centre (Université catholique de Louvain). BF2i laboratory was specifically supported by INSA Lyon and INRAE. The funders had no role in study design, data collection and analysis, decision to publish, or preparation of the manuscript.

## Notes

### Competing Interest Statement

The authors have declared no competing interest.

