## Supplementary figures and images for "The di-symbiotic systems in the aphids *Sipha maydis* and *Peryphillus lyropictus* provide a contrasting picture of recent co-obligate nutritional endosymbiosis in aphids"

### Figure S1.jpg

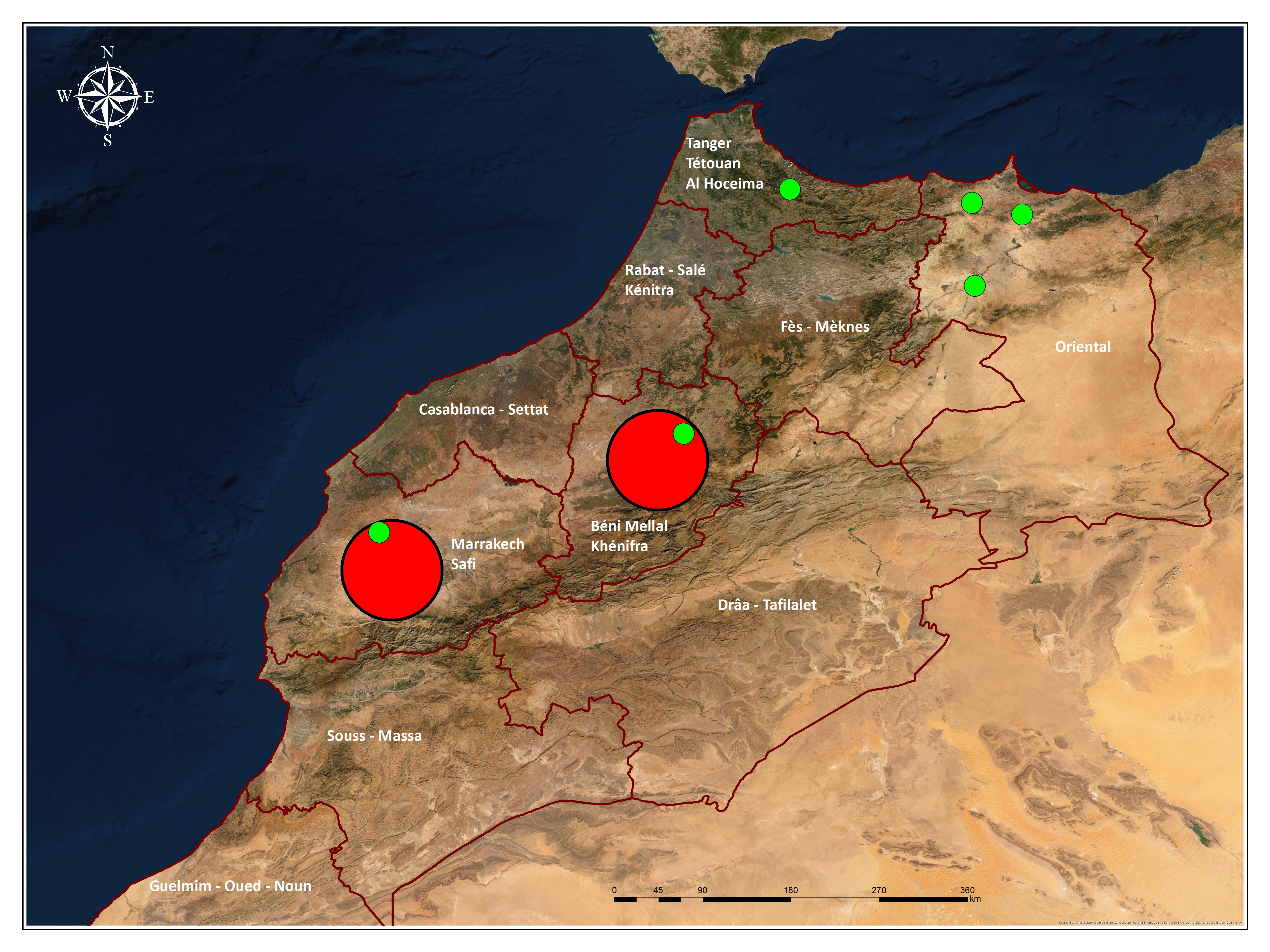
